# Demonstration of altruistic behaviour in rats

**DOI:** 10.1101/805481

**Authors:** Ushnik Das, Anshu Kumari, Shruthi Sharma, Laxmi T. Rao

## Abstract

A pilot study demonstrating complex cognitive abilities of altruism in rats by observing if rats would voluntarily put themselves in distress solely in order to help another conspecific irrespective of kinship and without any expectation of reward in terms of food or mate in return.

## Introduction

Haldane(1932) was one of the first men to describe altruistic traits as a trait which is “socially valuable but individually disadvantageous”. Since then, there have been several definitions and models describing altruism along with its plausible path of evolution as summarised by Kerr(2004). For the purpose of this study, the following definition of altruism by Uyenoyama(1980) has been adopted:

> “Altruism involves the sacrifice of fitness on the part of a set of individuals (the altruists) in order to increase the fitness of another set of individuals within the group, where fitness is defined for a set of individuals as the average number of offspring surviving to reproduce per individual in that set.”

There have been innumerable sightings of altruistic behaviour in the animal kingdom, ranging from dolphins saving swimmers from predating orca whales to dogs adopting kittens and other stray animals. But, there is only a handful of empirical evidence or scientific literature where altruism has been observed as per the definition concerned in this study. Sherman(1985) and Seyfarth (2012) reported that Belding’s squirrel and Vervet monkeys gave out alarm calls when sensing a predator, which gave out the caller’s precise location to the predator but helped it’s kin in escaping and thus, surviving. The latter also reported that the monkeys were more likely to produce alarm calls in the presence of close kin than rivals/strangers. Hare(2010) reported yet another example of altruistic behaviour where Bonobos voluntarily shared a highly preferred food item with another even though it had the opportunity of consuming it alone. Regurgitating and sharing of food by well-fed vampire bats to feed starved conspecific irrespective of kinship or bonding was observed by Gerald(2013). In the marine world, Pitman(2016) observed humpback whales interfering when mammal-eating killer whales attack other species. The humpbacks on several occasions risked its own well-being while saving the victim animals, even of different species.

Scientists like Marco(2012), Stevens(2004), Penn(2007) and Wall(2010) have supported the fact that non-primates such as rats don’t have the cognitive capabilities required to demonstrate the complex emotions of altruism or even empathy [Wrighten(2016)]. But there have been empirical evidences which point to the direction that these rodents might be able to showcase altruistic traits in themselves. Rice(1962) was the first person to report altruism (though not as per our definition: devoid of “sacrificing it’s own fitness”) in rats which had followed heavy criticisms from Lavery(1963). Later Bartal(2011) demonstrated that rats would free another trapped rat and share food with it voluntarily. This sharing of a highly palatable food item could be thought of as an altruistic behaviour. Dolivo(2015) showed that the fact that a rat would help another or not was determined by not only on whether it had received help from the other but also on the quality of help(in terms of highly/lowly preferred food) it received earlier. Though superficially, it may seem that experiments by Bartal and Dolivo are examples of altruism but not so on a deeper analysis. The rats in the former experiment weren’t starving and hence sharing of food item wouldn’t have decreased their fitness. Similarly, the helper rat in Dolivo’s experiment isn’t affecting it’s fitness in any way to help the other. Hence as per our definition of altruism, these two experiments fall short. What would be interesting is to see if a starved helper rat, goes out of his way to release a trapped rat and then share food with it. Or the case where a well-fed rat would acquire food for a starved one in case of Dolivo’s experiment. Therefore there hasn’t been a study where a rat would help another at the cost of its own fitness. So here, we’ve designed an experiment to check just that. Whether or not a rat would put itself in discomfort and help another without any expectation of reward in terms of food or mate.

Sutanto(1994) reported that stressful conditions in rats results in various pathological/biochemical vulnerabilities affecting it’s own fitness while Bradesi(2005) reported that rats would actively avoid water wells (8cm in his experiment) which results in chronic stresslike conditions. So on the basis of the last two reports one can say that rats would try to avoid getting into water as it is a stressful environment to them.

Now, according to the definition of altruism, if the rat voluntarily puts itself in distress(for instance, crossing a water well) hence reducing its own fitness just to rescue its conspecific, then it can be well defined as an act of altruism.

### Aim

To show that rats have altruistic traits in themselves.

### Hypothesis

Rats would voluntarily attempt to put itself under discomfort or stress with the sole purpose of helping/rescuing another rat who is in trouble.

Thus, our research would attempt to prove the mentioned hypothesis through an experimental setup to show altruism in rats.

## Methodology

### Subjects

The present study is an extension of the ongoing PhD work (Kumari Anshu) in the lab.

Male Wistar rats and female SD rats of 4–5 months were used for the present study. These rats were housed 2 per cage in Central Animal Research Facility (CARF), NIMHANS, Bengaluru in a climate-controlled room having light–dark cycle with food and water available ad libitum.

### Experimental apparatus

#### Empathy training and Altruism test

The apparatus consisted of the restraining tube previously used by Bartal(2011) and 3 rectangular boxes each of dimensions 45×30×30 cm interconnected via a large gate. The first and the third box had an elevated basement of ∼9cm.

#### Anxiety test

The apparatus (Coulbourn Instruments, USA) consisted of light and dark compartments, each 26 × 26 cm and connected by 8 × 8 cm guillotine doorway. The light chamber was illuminated with a bright light (∼300 lx) on the ceiling. The door was programmed to open at the 5th second immediately after initiating the protocol.

### Experimental protocol

The experiment can be divided into 3 stages:

Stage 1: Habituation and water avoidance test

Stage 2: Training to test empathy behaviour in rats

Stage 3: Altruism test

#### Stage 1: Habituation and water avoidance test

The rats were exposed to the experimental 3-chambered apparatus thrice a day for 10mins each. After 2 days of exposure,the mid chamber was filled with water till the level of the water surface was same as the elevated bases of the other two boxes. Each rat was individually placed in the first box (box A) and it was observed for 15 mins whether the rat would cross the water well or not.

#### Stage 2: Testing procedure for checking empathy behaviour in rats

The protocol of Bartal(2001) was followed. The empathy chamber with a trapped rat (TR) and a free rat(FR) was kept in the third box (box C) with the gate to the other box closed. The timer was started and the setup was left undisturbed for 10 mins. After 10mins, if the FR was unsuccessful in opening the chamber, the experimenter would open the chamber gate by 20-30% and leave the setup undisturbed. After 5 mins, if the rats were still unsuccessful in opening the chamber, the experimenter would completely open it and leave the subjects to explore the box for another 5 mins. This was done till the point the FR understood how to open the gate by itself. If the FR opened the gate before the experimenter, both the rats were left undisturbed for 5 mins before putting them back into their home cages.

4 male and 4 female rats were randomly chosen for the experiment. A trial test was done to check each rat’s activity as a FR. 2 of most active rats from male(M_1_-M_2_) and female(F_1_-F_2_) each were chosen as the selected FRs.

They were exposed to the said experiment till they successfully opened the chamber for 3 consecutive days. After 5-7 successful sessions, the empathy chamber was then kept in box C with a TR, while a FR was kept in box A separated by the dry second box(B) and the experiment was continued at a rate of 2 sessions per day, till the rat had 32 sessions in total. The time taken to open the entrapment chamber in each case was recorded.

#### Stage 3: Altruism test

Box B was filled with water upto the base level of the first and third box. A FR was kept in box A and a TR in box C separated by the water well. The latency in the time taken to come in contact with water and finally cross the well was recorded along with the time taken to free the TR.

Every FR was experimented with a familiar(cagemate) TR followed by an unfamiliar TR for 10 sessions each.

#### Anxiety test

Each rat was individually placed in the dark environment and allowed to explore freely in both light and dark environments. The anxiety test was run for 5.5 mins for each rat. Five second after the placement in the dark chamber, door to the lit chamber opens. The time spent in the brightly lit environment, latency of first transition and the number of transitions between the compartments were quantified offline. The parameters -- time spent in dark chamber, latency of first transition and number of transitions served as measures of anxiety behaviour.

*After every session in each stage, the apparatus was thoroughly cleaned with 70% alcohol.

## Results

### Stage 1: Habituation session

Total of 4 rats were used to evaluate the altruistic behaviour in rats. Out of which only 3 rats were successful in completing all the three steps of the experiment. At the first step during habituation stage, all rats explored all the three chambers -- A,B and chamber C. All four rats exhibited exploratory locomotion, rearing and grooming behaviour during 10 minute sessions. On day 3, when the chamber B was filled with water, none of these 4 rats stepped into chamber B filled with water and restricted themselves to exploring only the first chamber A indicating that rats exhibit aversive behaviour to the water.

### Stage 2: Empathy behaviour

As shown in Fig.(2.a), all the 4 rats were successful in releasing the trapped rats by session 16, but significant difference was first observed in session 19(p=0.0323) with a latency of less than 90 seconds (mean latency of 43.5s and SEM of 15.25s). All subjects showed a decrease in the time taken to free the trapped rat with increasing sessions, indicating progressive learning as evidenced by F_(20,60)_= 3.957, p<0.0001. Individual rats took different number of sessions to learn to open the chamber door varying from session 6 to session 16 which in turn varied from 152 secs for M_1_ to 374 secs for M_2_. But after the end of 28 sessions, the time taken by all the 4 rats converged to have a mean of 14.6 seconds with a SEM of just 2.36 secs. Repeated one-way ANOVA using Bonferroni’s multiple comparison test was conducted to observe a statistically significant P value of <0.05 when session 13 (chosen because by then majority of rats had learnt to open the door) was compared with session 28 and onwards, indicating a significant drop in latency due to learnt behaviour. There was also a significant difference between learning abilities of different rats as is evident from F_(3,60)_=13.01, p<0.0001.

**Fig (1):**
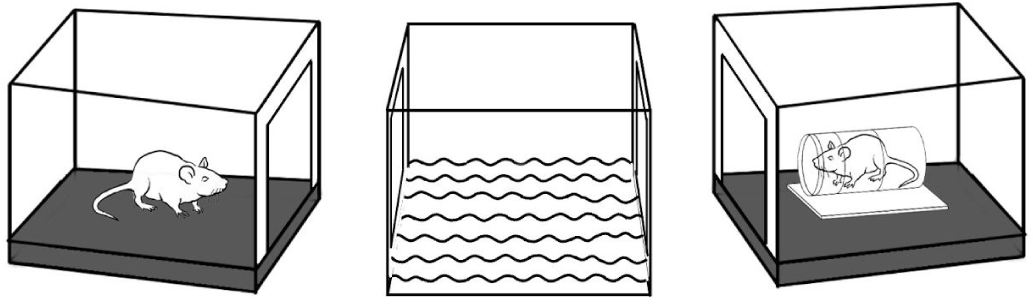
The experimental apparatus with the central one having water. Boxes are shown separated for clarity.

**Fig (2):**
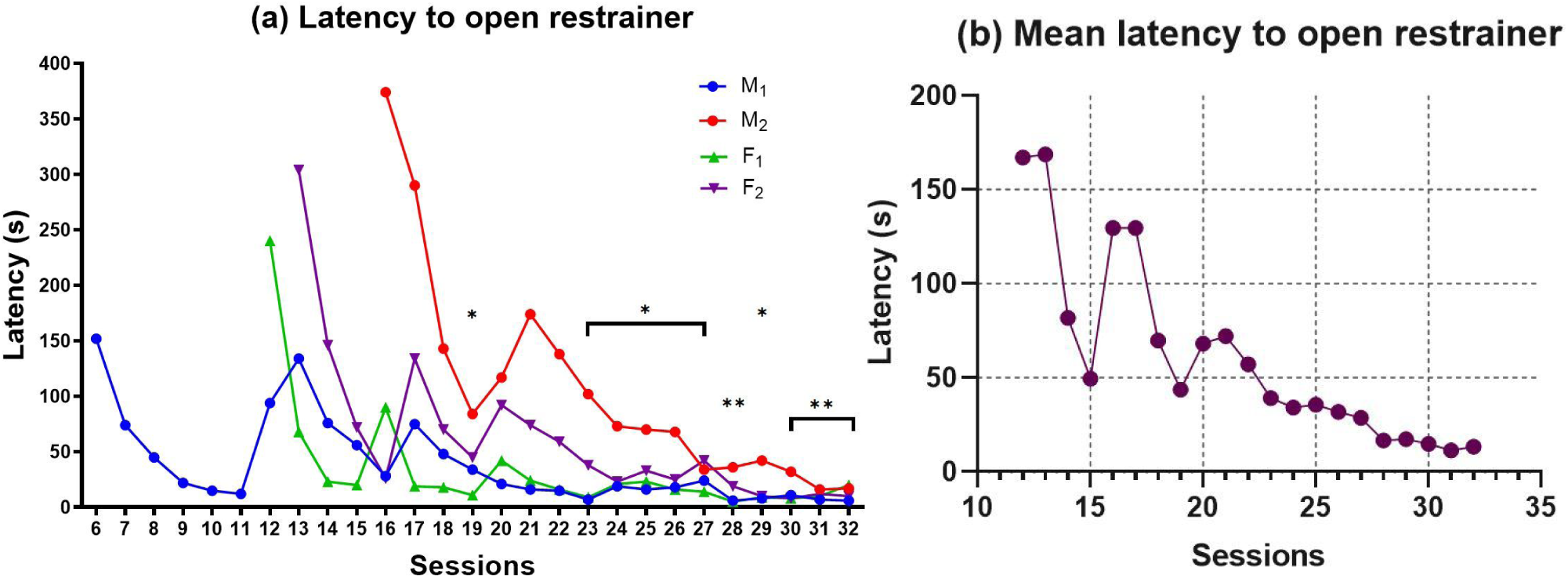
(<u>a</u>)Individual and (<u>b</u>)Mean latency of the rats in opening the restraining chamber where, * and ** denotes a p value of <0.05 and <0.01 respectively, when compared to session 13

### Stage 3: Altruism behaviour

3 out of 4 rats crossed the water barrier in all the sessions carried out, the remaining one rat only crossed twice out of 20 trials and hence has been kept out of consideration.

As seen in Fig.(3), when first exposed to the water barrier, all the 3 rats crossed it with a significantly decreasing value in latency (F_(9,18)_=3.68, p=0.009). There was also a significant difference observed amongst individual rats in learning to cross the water barrier (F_(2,18)_=9.228, p=0.0017).

**Fig.(3):**
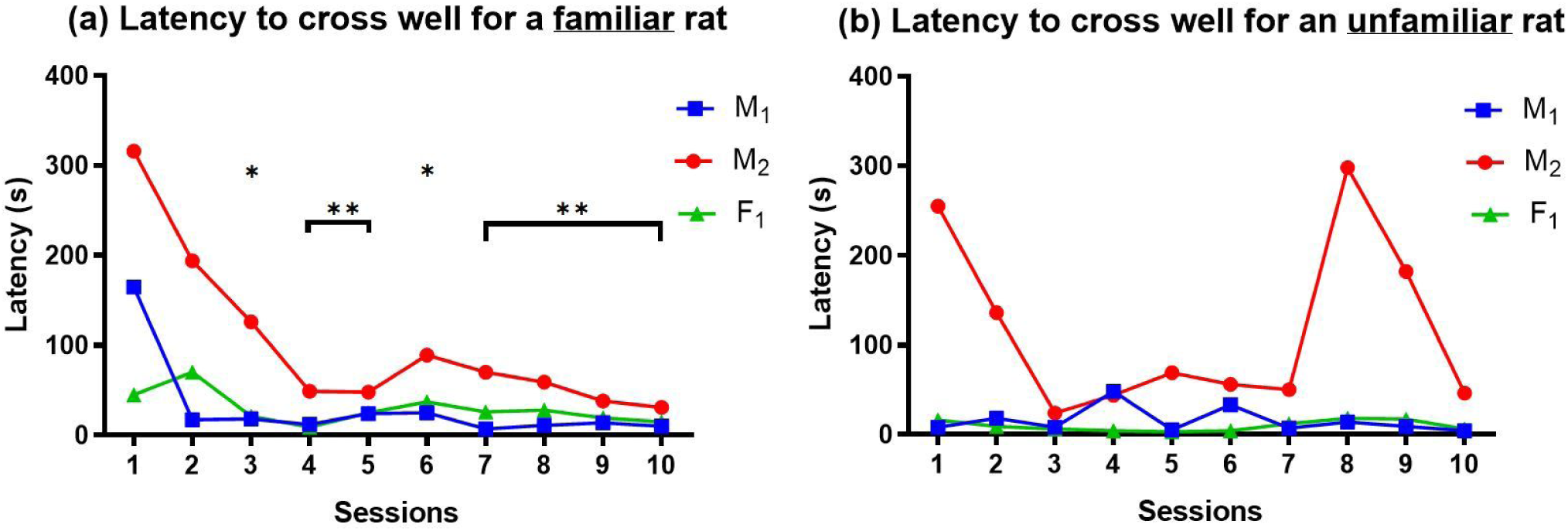
Latency of crossing the water well with a trapped (<u>a</u>) familiar and (<u>b</u>) unfamiliar conspecific where, * and ** denotes a p value of <0.05 and <0.008 respectively, when compared to session 1

After 10 sessions each, when these trained rats were exposed to trapped unfamiliar conspecifics there wasn’t any significant trend observed in their behaviour towards increasing or decreasing latency for crossing the water barrier (F_(9,18)_=1.054, p=0.4385).

According to Fig.(4), once the subjects cross the barrier, no significant difference in the latency to open the chamber for familiar (F_(9,18)_=0.9298, p =0.5233) or unfamiliar (F_(9,18)_=2.378, p=0.0562) rats is observed. But the mean time(±SEM) to open the cages for familiar and unfamiliar rats were observed to be 15.03(±4.76) seconds and 7.67(±2.31) seconds respectively.

**Fig.(4):**
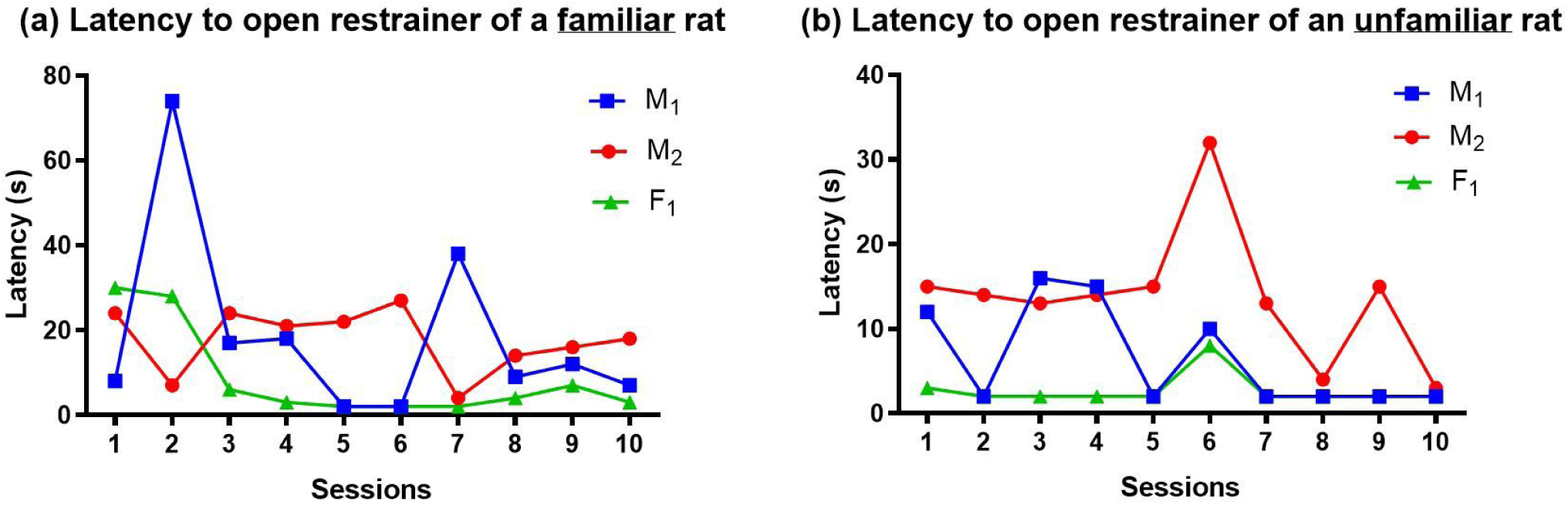
Latency of opening the restrainer after crossing the water well with a trapped (<u>a</u>) familiar and (<u>b</u>) unfamiliar conspecific

where, * and ** denotes a p value of <0.03 and <0.008 respectively, when compared to session 1 From Fig.(5.a), we observe that there is a decreasing trend in latency to cross the water barrier, statistically evident by F_(19,38)_=1.968, p = 0.0374. The graph shows the significant differences observed when those sessions were compared to session 1, as a result of the learning behaviour of traversing a new challenging path. After 10 sessions with a familiar TR, when it was experimented with an unfamiliar TR, there wasn’t any statistically significant difference observed when compared with session 11, as the rats had already learnt how to traverse the water barrier. Since opening of the restrainer is a learnt behaviour, there isn’t any trend observed in Fig.(5.b) nor is there a significant difference in the latency of freeing the trapped rat for familiar and unfamiliar conspecifics, as evident by F_(19,38)_=1.370, p =0.1998.

**Fig.(5):**
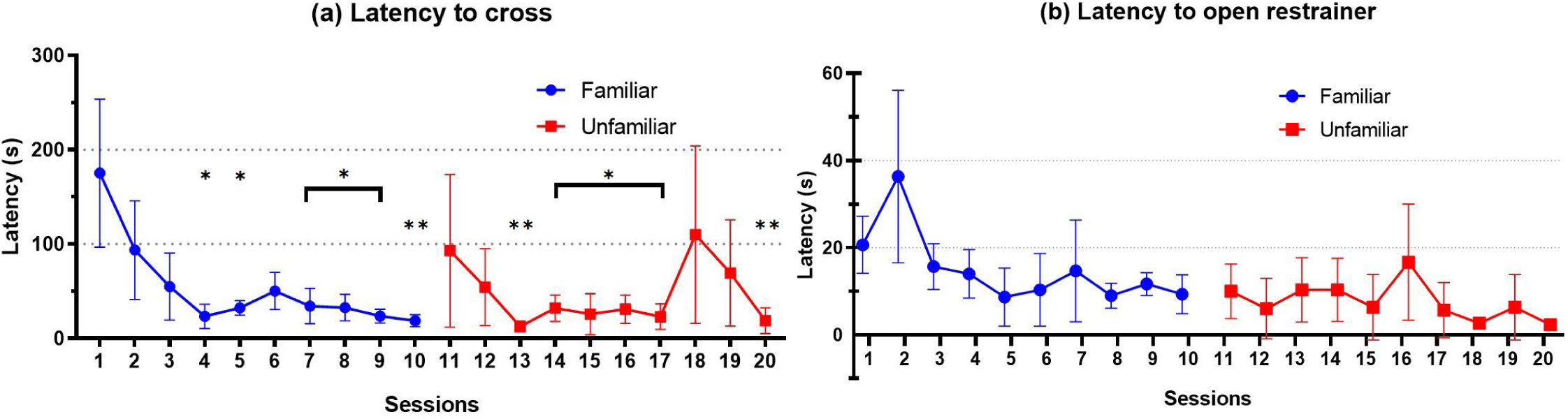
Consolidated latency of (<u>a</u>) crossing the water barrier in chamber B and (<u>b</u>)opening the restrainer in chamber C

### Anxiety test

From Fig.(6), 3 out of 4 subjects showed similar results in the light-dark test with a mean time of 46.33±7.88 seconds spent in the light chamber while the other subject(M_2_) spent a considerably low duration of 17 seconds. The 3 former rats traversed through the gate on an average of 4.33±0.88 times in the total duration of 5.5mins while the latter one just had 2 crosses through the gate. The latency of the first transition between chambers had similar values in all the 4 rats with a mean of 5.5±0.96 seconds.

**Fig.(6):**
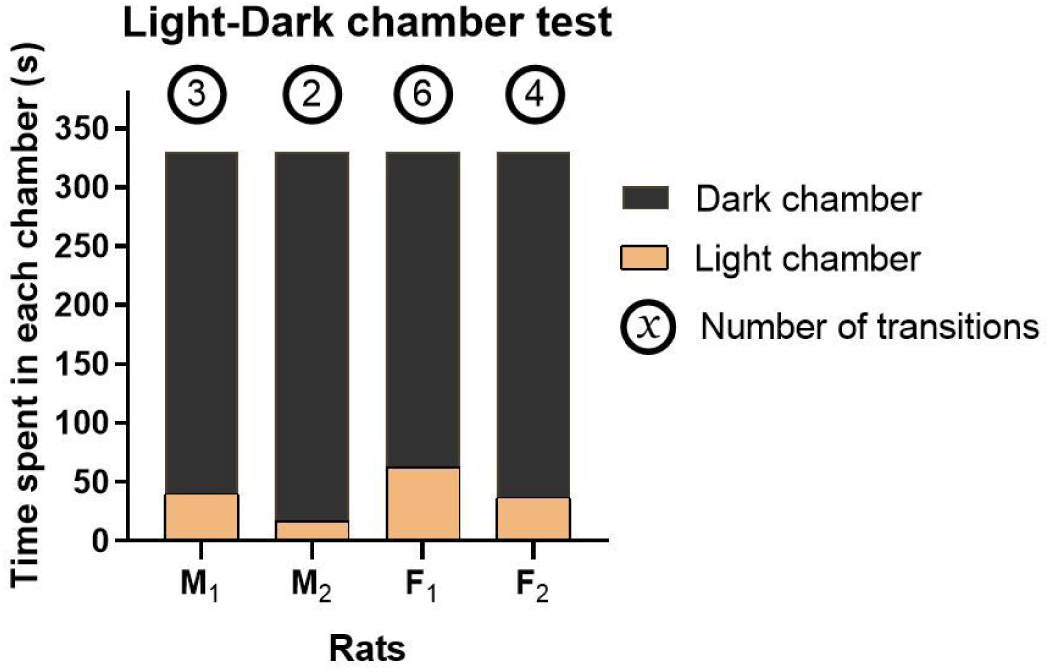
Individual time duration spent in the Light-Dark chamber along with total transitions

## Discussion

Our hypothesis was that if the rats put themselves in stress, here crossing the water well, just to rescue its conspecific then it is an act of altruism. Through our experimental study we confirm that the rats would not only cross the water barrier to save its distressed cagemate(familiar rat) but also to rescue a conspecific which it had no prior contact or relation with(unfamiliar rat).

Even though the rats were familiar with the 3 chambered apparatus due to repeated exposures prior to the control test, none of the rats ventured into the water well or tried to cross it by any means to reach chamber C. This is a testimony to the report of Bradesi(2005) that water wells are indeed a stressful condition to rats and thus it makes sense that they would avoid it whenever possible as it would lead to a decrease of their own fitness as pointed out by Sutanto(1994).

In the empathy chamber training sessions, the first-hand motivation to attempt to open the chamber is due to an empathetic response felt by the free rat after it realises that it’s conspecific is in distress [Bartal(2011)]. Through the graphs one can observe that different rats took different amounts of time to understand how to open the gate. This is attributed to the fact that each rat has different learning abilities. But nonetheless, all the rats showed a progressively decreasing trend in the time taken to free the trapped rat which is again a testimony to the learnt behaviour in rats. Since all the rats opened the gates within a minute in the last 5 sessions as compared to none in the first 5, it is evident that there is a statistically significant difference here. So, the ANOVA test conducted was used to compare the 13^th^ session with the latter ones to check if there is any significant trend in the latency. The 13^th^ session was chosen because 3/4 rats had learnt to open the gates by then and hence any trend observed henceforth would dictate the trend in the majority of the population. As it was seen through the test, there was a significant difference not only in the total time taken to open the gates in between sessions but also amongst individuals. Both of which supports the latency of learning and its individualistic variations in rats.

At the end of the empathy training sessions the 4 rats were able to open the chamber in almost the same time. Now they were introduced to the water barrier. As one observes through the graphs, the latency to cross shot up as it was a new challenge for the rats. But with subsequent exposures, they slowly got used to the new stressful condition and also learnt to cross it: either by jumping or by swimming through it rapidly. The decreasing trend in the latency and a statistically significant difference between the first and subsequent sessions are again a testimony to the trend observed in learnt behaviour.

This sole act of putting oneself in this stressful environment of water is considered altruistic. If we compare it to the control observation, the same group of rats completely avoided the water body but after it realised the presence of a trapped conspecific in need of its help, it tried its level best to come across and help the trapped one. What was interesting to observe was that when an untrained rat (i.e. a rat which hadn’t experienced empathy training sessions) was kept in box A with a trapped conspecific, it would spend more time at the water edges which could be possibly due to its own responses to the distress call of the trapped rat. But never did it once try to cross the water barrier. One can conclude from this observation that the motivation to help the trapped conspecific increased after it was sure of the fact that it could help the trapped one from its distress. After 10 sessions of trapped familiar conspecific, the rats were tried with trapped unfamiliar conspecific. One observed a slight increase in the mean latency of crossing, but this is attributed to the experimental protocol which had a gap of 2 days between the familiar-unfamiliar sessions, so it is possible that one of the rats might have taken some time to recall the learnt behaviour.

The re-learning was quick as we can see that within 3 sessions, the rats began to perform as they did for the familiar conspecifics. A similar trend is observed in the time taken to free the trapped rat after it reaches box C. It is also noteworthy to mention that the behaviour of the rat doesn’t change with a familiar or unfamiliar trapped rat which shows that this is indeed an example of altruism and not kin selection.

On observing the anxiety data, we couldn’t find a strong relation between it and the trend observed in other experiments here. For instance, the rat F_2_: though it showed similar values to M_1_ and F_1_ in light-dark test but it crossed the barrier only twice out of 20 sessions. Since it performed well in the empathy training test, one can’t say it’s a poor learner and didn’t know that a trapped rat was on the other side. The two plausible explanations of not crossing the barrier could be perhaps that it wasn’t as anxious to brightly lit environment as it was to water wells or perhaps it wasn’t as altruistic as the other subjects. Since it is an experiment of anthropomorphism, the latter explanation seems equally valid as far as the observations are concerned. But one should also note that even though M_2_ crossed the water body every time to help its conspecific, it was the most anxious subject in the light-dark test with the least number of transitions and duration in light chamber. Added to this, it had the most inconsistent performance in latency of crossing as well as latency to free the trapped rat. After the gap of 2 days between familiar-unfamiliar rat session, M_2_ had taken an unusually long time to cross the barrier as compared to the other subjects, but it did relearn in subsequent sessions. One can conclude from this observation that even though M_2_ was most anxious towards exposure to light but it wasn’t so much anxious with it’s exposure to water added with the altruistic motivations.

It is also noteworthy to mention that our experiment was conducted with a small sample space consisting of just 4 rats out of which 3 had showcased clear signs of altruism. Now, with a larger sample space one can further reaffirm this result to make it a statistically significant claim. Till date, there hasn’t been any empirical evidence of altruism in non-primates hence such a result would mark as a milestone in animal behavioral studies and our understanding of human cognition and anthropomorphism in the animal kingdom. If we look at the future research prospective, there is an unimaginable scope of research if such a claim proves to be correct, especially in the fields of neurobehavioral disorders such as autism where altruistic traits are remarked to be absent in the affected patients.

## Conclusion

Through this experimental setup, we were able to prove the hypothesis by demonstrating how rats would voluntarily opt to put themselves in distress in order to rescue a troubled conspecific, irrespective of kinship and without any expectation of a reward in terms of food or mate. This perfectly aligns with the definition of altruism adopted for this study and hence: Rats can be regarded as organisms capable of showcasing complex cognitive features of altruistic behaviour.

